# CRISPR-Cas systems in *Fusobacterium*: An untapped genetic frontier

**DOI:** 10.1101/2022.01.21.477265

**Authors:** Ariana Umaña, Daniel J. Slade

## Abstract

*Fusobacterium nucleatum* has recently received significant attention for its strong connection with the acceleration and gravity of multiple cancers (e.g., colorectal, pancreatic, esophageal). However, our understanding of the molecular mechanisms that drive infection by this ‘oncomicrobe’ have been hindered by a lack of universal genetic tools. Herein we report a global bioinformatic identification and characterization of multiple *Fusobacterium* CRISPR-Cas adaptive immune systems including Cas13c, and detailed report of the proteins, spacer/repeat loci, *trans-* activating CRISPR RNA (tracrRNA), and CRISPR RNA (cRNA) from a Type II-A CRISPR-Cas9 system. Since most *Fusobacterium* are currently genetically intractable, this CRISPR-Cas bioinformatic roadmap could be used to build new genome editing and transcriptional tuning tools to characterize an increasingly important genus of human opportunistic-pathogens connected to the onset, progression, and severity of cancer.

## INTRODUCTION

Fusobacteria are anaerobic Gram-negative bacteria, with the most well studied human and livestock opportunistic pathogens of this genus being *F. nucleatum and F. necrophorum*. Due to robust restriction modification systems that hinder genome manipulation, *Fusobacterium* genetics have classically been considered nearly impossible, resulting in few bacterial strains that can be genetically modified for detailed mechanistic studies. Despite recent advances in *F. nucleatum* genetic tools, mainly in the strain *F. nucleatum* subsp. *nucleatum* ATCC 23726, all other species outside of *F. nucleatum* have nearly fallen into the category of genetically intractable ^1-3^, with the exception being a recent study highlighting a conjugation method for genetic manipulation of *F. necrophorum* ^*4*^.

Recent advances in our understanding of Clustered regularly interspaced short palindromic repeats-CRISPR-associated proteins (CRISPR-Cas) systems have been exploited by molecular biologists to allow for efficient and robust genome editing in both prokaryotic and eukaryotic systems ^5-7^. CRISPR and Cas proteins comprise the basis of adaptive immunity in bacteria and archaea ^8^. CRISPR-Cas systems identify and cleave non-host DNA and RNA (viruses and plasmids), and deposit a ∼30 bp segment into its host genome at defined CRISPR loci (host DNA repeats and foreign protospacer, or ‘spacer’ DNA) to provide organism- and sequence-specific defenses ^9^. During repeat infections, CRISPR-Cas complexes recognize these specific sequences and rapidly clear the invading DNA through active nucleotide cleavage ^10^. CRISPR systems have been characterized in 87% of archaea and 47% of bacteria and have been proposed to proliferate through horizontal gene transfer across bacteria as a basis of adaptive immunity ^9^.

CRISPR arrays are divided into three main categories and 16 distinct subcategories based on the *cas* genes encoded within bacteria and archaea ^11^. However, the majority of genome editing studies have used the Cas9 protein from *S. pyogenes* ^11^. While the use of *S. pyogenes* Cas9 in prokaryotic and eukaryotic systems has been achieved, our goal was to characterize *Fusobacterium* CRISPR-Cas proteins to determine if these proteins will be more efficient for future gene modulation studies in an AT rich genus of bacteria. The discovery and characterization of native Class 2 effectors in *Fusobacterium* will potentially enhance the application and development of CRISPR systems for genome engineering. Herein, we report a comprehensive bioinformatic characterization of multiple *Fusobacterium* CRISPR-Cas adaptive immune systems, focusing on Cas9 and the newly described Cas13c present in *F. necrophorum* subsp. *funduliforme* 1_1_36S. This data, coupled with a recent study characterizing an active CRISPR system in the strain *F. nucleatum* subspecies *nucleatum* ATCC 25586 ^12^, provides hope that the development of new CRISPR-based genetic tools will enhance our understanding of fusobacterial physiology and pathogenesis.

## MATERIALS AND METHODS

### Use of genomic information for bioinformatic analysis and identification of CRISPR-Cas systems in eight *Fusobacterium* strains

*Fusobacterium* genomes (*F. nucleatum* subsp. *nucleatum* ATCC 23726 (GCA_003019785.1), *F. nucleatum* subsp. *nucleatum* ATCC 25586 (GCA_003019295.1), *F. necrophorum* subsp. *funduliforme* 1_1_36S (GCA_003019715.1), *F. varium 27725* (GCA_003019655.1), *F. ulcerans 49185* (GCA_003019675.1), *F. mortiferum* 9817 (GCA_003019315.1), *F. gonidiaformans* 25563 (GCA_003019695.1), and *F. periodonticum* 2_1_31 (GCA_003019755.1) were used to extract all sequences to analyze using the CRISPROne web server ^13^ and CRISPRCasFinder ^14^. CRISPROne and CRISPRCasFinder were used to predict all CRISPR-Cas associated elements as well as the repeat arrays.

### Bioinformatic analysis of CRISPR-Cas Class 2 systems and identification of effector domain

The genome sequence from *F. necrophorum* subsp. *funduliforme* 1_1_36S (GCA_003019715.1) was used to predict the open reading frame for the CRISPR-Cas systems. An open reading frame of 1337 amino acids for Cas9 and 1121 amino acids for Cas13c was identified using the web server pHMMER ^15^ and stand-alone HMMER software package, respectively.

### Phylogenetic Tree construction

Full CRISPR-Cas phylogenetic analysis was created using the micropan ^16^ plugin in Rstudio by using only CRISPR associated proteins to build the tree. The gene maps in Figure 1B were created using the gggenes ^17^ plugin in Rstudio. Additional phylogenetic trees for Cas9 and Cas13c were created in Geneious 9.18 utilizing the multiple sequence alignment from SMARTBLAST and adapted in Affinity Designer.

**Figure 1.**
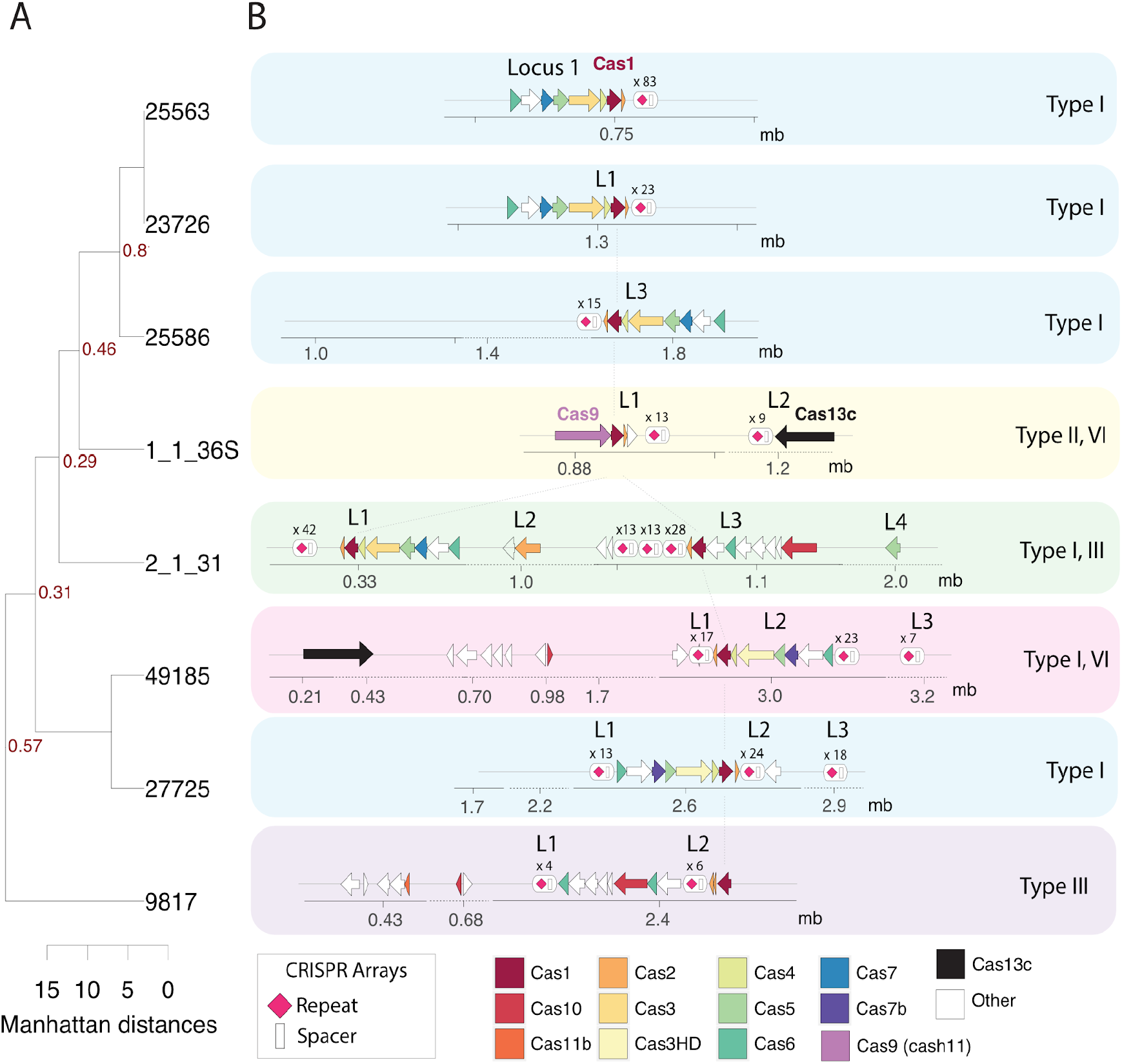
CRISPR-Cas systems in *Fusobacterium*. **A**. CRISPR-Cas phylogenetic tree of eight *Fusobacterium* strains. **B**. A total of eight *Fusobacterium* strains were utilized for bioinformatic analysis and identification of CRISPR-Cas systems. A majority of the identified CRISPR-Cas systems in *Fusobacterium* belong to Class 1 and only *F. necrophorum* susbp. *funduliforme* 1_1_36S and *F. ulcerans* 49185 encode for Class 2 systems.

### Structure Prediction

The structure prediction of Cas9 from *Fusobacterium necrophorum* subsp. *necrophorum* 1_1_36S was performed in the web-based suite Phyre2 ^18^. CRISPR RNA structure prediction was done using NUPACK ^19^.

## RESULTS

### Comparative Analysis of *Fusobacterium* CRISPR-Cas systems

A total of eight *Fusobacterium* genome sequences were utilized for bioinformatic comparative analysis and the composition, occurrence, and diversity of CRISPR-Cas systems. Presently, there are two classes and six types of CRISPR-Cas systems based in the organization of their effector modules. A majority of the identified CRISPR-Cas systems in *Fusobacterium* belong to Class 1 (77% of CRISPR-Cas systems in the eight genomes) and interestingly, only *F. necrophorum* susbp. *funduliforme* 1_1_36S and *F. ulcerans* 49185 encode for Class 2 systems (23% of CRISPR-Cas systems in the eight genomes). The overall analysis of CRISPR-Cas systems is shown in **Table S1**.

To depict the correlation among *Fusobacterium* species and strains, a CRISPR-Cas system-based phylogenetic tree was constructed. As shown in **Figure 1A**, the eight *Fusobacterium* strains are divided into five distinct clades according to their predicted CRISPR-Cas systems. Interestingly, *F. nucleatum* ATCC 23726 (Lineage 1) clades with *F. gonidiaformans* 25563 (Lineage 3) and *F. periodonticum* 2_1_31 (Lineage 1) clades with *F. necrophorum* subsp. *funduliforme* 1_1_36S (Lineage 3).

The rapid evolution of CRISPR systems explains the diversity of systems from highly similar strains. For instance, *F. nucleatum* ATCC 23726 and *F. nucleatum* ATCC 25586 are the closest phylogenetically when using whole genome phylogenetic analysis, yet as seen in **Figure 1A**, they are far more divergent in their CRISPR loci. Additionally, the analysis revealed that *F. mortiferum* 9817 CRISPR-Cas systems have less conservation when compared to the other *Fusobacterium* strains.

Estimation of spacer numbers and sequences among *Fusobacterium* species exhibited substantial diversity with 0% (0/348) of the spacers shared by eight strains. As shown in **Table S1**, the quantity of predicted spacers varies among the strains, ranging from 4 to 83, demonstrating the biological heterogeneity of *Fusobacterium* strains that encode an adaptive defense mechanism. Type I-B presented a higher number of spacers when compared to the other CRISPR-Cas system types.

### Classification of CRISPR-Cas systems in *Fusobacterium*

To determine the classification and assortment of CRISPR-Cas systems across the eight *Fusobacterium* strains, we utilized the CRISPRCasFinder and CRISPROne web server that enabled the discovery of direct repeat consensus sequences within a genome as well as the characterization of related spacers and *cas* genes ^13,14^. This analysis detected a total of 13 CRISPR-Cas systems belonging to both Class 1 (Type I, Type III and Type IV,) and Class 2 (Type II, Type VI): six Type I (54.54%), one Type II (9.09%), two Type III (18.18%), and two Type VI (18.18%) while no Type IV or V was identified. Therefore, we conclude that Type I is the predominant CRISPR-Cas system among the studied *Fusobacterium* strains. Detailed illustrations of the CRISPR-Cas systems identified among the strains analyzed in this study are summarized in **Figure 1B**. *F. nucleatum* ATCC 23726, *F. nucleatum* ATCC 25586, *F. gonidiaformans* 25563 and *F. periodonticum* 2_1_31 Type I-B is a Class 1 CRISPR-Cas loci that includes full gene sets to achieve adaptive immunity: *cas1, cas2* and *cas4* for spacer integration or adaptation, *cas6* for crRNA-processing and *cas3, cas5, cas7*, and *cas8* for interference. Both *F. ulcerans* 49185 and *F. varium* 27725 include a nearly complete Type I loci, but they do not encode for the *cas8* gene. Additionally, most CRISPRs of Type I-B CRISPR-Cas systems discovered in this analysis were encoded downstream of the *cas2* gene and/or upstream of the *cas6* gene. Worth noting, *F. nucleatum* ATCC 23726 and *F. gonidiaformans* 25563 presented an inverted loci composition of the Type I-B CRISPR-Cas systems identified **(Figure 1B)**.

Furthermore, in *F. periodonticum* 2_1_31, Type I and Type III systems appear to combine as they often coexist ^20^. Type III systems have been divided into four major subtypes (III-A to D) 21,22. A complete operon of *cas* genes associated with adaptation, crRNA-processing and interference steps was identified in *F. periodonticum* 2_1_31. Additionally, Type III systems are found to frequently encode for accessory genes near the core *cas* gene cluster.

### Analysis of a *F. necrophorum* Type II CRISPR-Cas system and identification of putative repeats/spacers/PAM sequence

The only Type II-A CRISPR-Cas system identified in our analyzed *Fusobacterium* strains belonged to *F. necrophorum* subsp. *funduliforme* 1_1_36S. Type II-A is a Class 2 system that contains four adaptation genes; *cas1, cas2, csn2*, and *cas9* (**Figure 1**), where Cas9 acts as a nuclease responsible for invader DNA cleavage **(Figure 2)** ^23^.

**Figure 2.**
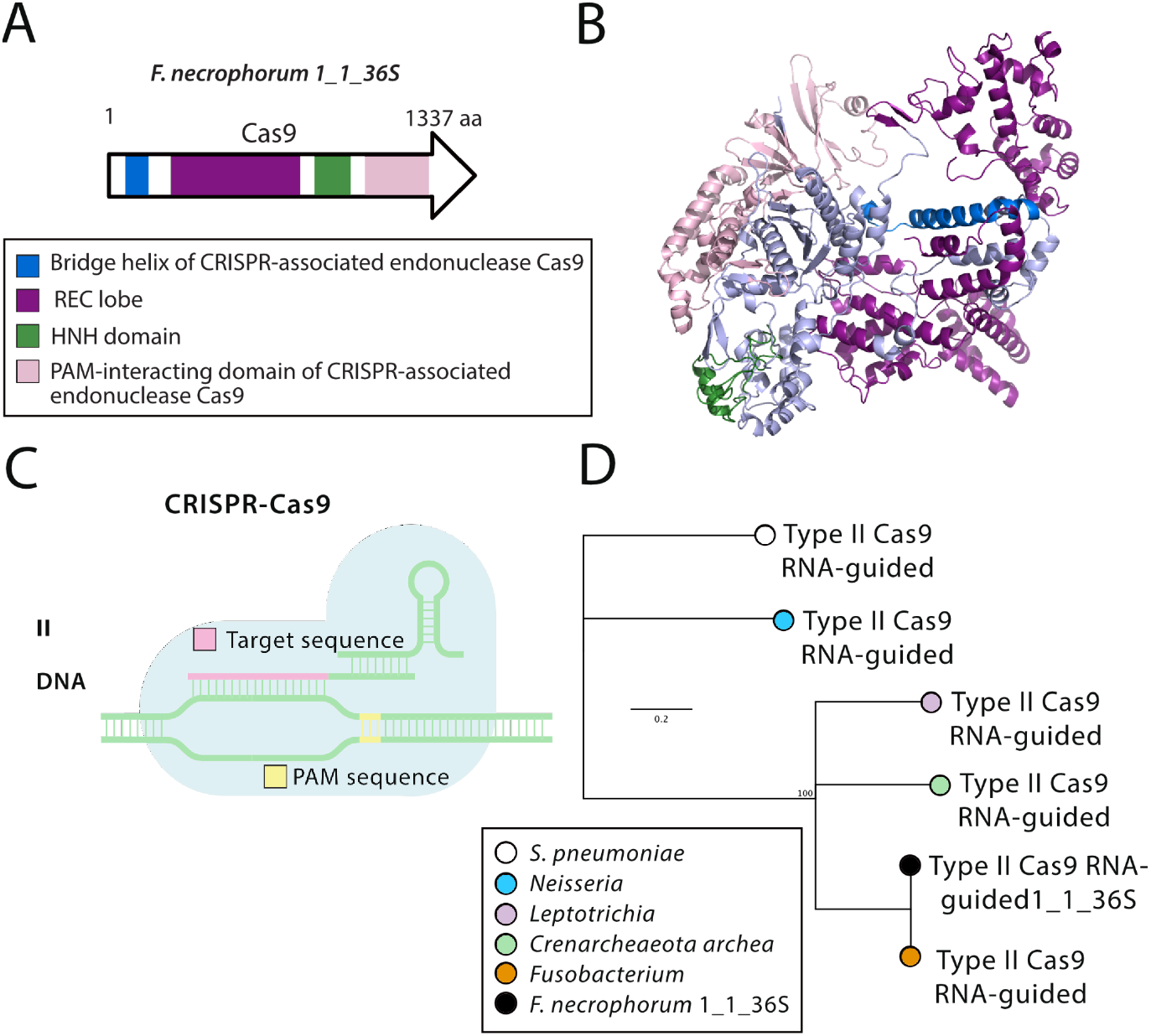
Class II CRISPR-Cas9 system in F. *necrophorum* subsp. *funduliforme* 1_1_36S. **A**. Effector domain organization in Cas9 with nuclease domains are detailed. **B**. Predicted structure of Cas9 with domains color coordinated with section A. **C**. Schematic of CRISPR-Cas9 in *F. necrophorum* subsp. *funduliforme* 1_1_36S. **D**. SMART-BLAST phylogenetic analysis of Cas9.

The Type II CRISPR-Cas systems function through the assembly of an effector module consisting of a single Cas9 protein, a crRNA and a tracrRNA. Cas9 is 1337 amino acids in size. A previous genome assembly of *F. necrophorum subsp. funduliforme* 1_1_36S had this protein as 329 amino acids, proving that complete and accurate genome sequences found in FusoPortal were key for identifying the proper open reading frames of these proteins ^24^. The predicted structure of Cas9 **(Figure 2B)** resembles the characteristic bi-lobed architecture from previously characterized Cas9 proteins, with the RuvC and HNH domains for nucleic acid cleavage **(Figure 2A)**.

In addition to Cas9, recognition and targeting of DNA is dependent on the crRNA and tracrRNA complex ^25^. For this reason, we characterized the tracrRNA element of the Type II CRISPR-Cas system in *F. necrophorum* subsp. *funduliforme* 1_1_36S. The first nucleotides predicted in the nexus include adenine residues that are known to be highly conserved among tracrRNAs of Type II systems as shown in **Figure 3A** ^25^ and the bulge interacts through a specific side-chain with the Rec1 domain or lobe through specific side-chain ^26^. Both bulge and nexus in tracrRNA are required for DNA cleavage, and the spacer sequence detailed in **Figure 3B** will determine the location of Cas9 endonucleolytic cleavage ^25^. Finally, the stems interact with Cas9 mainly through sequence-independent interactions with the phosphate backbone ^25^.

**Figure 3.**
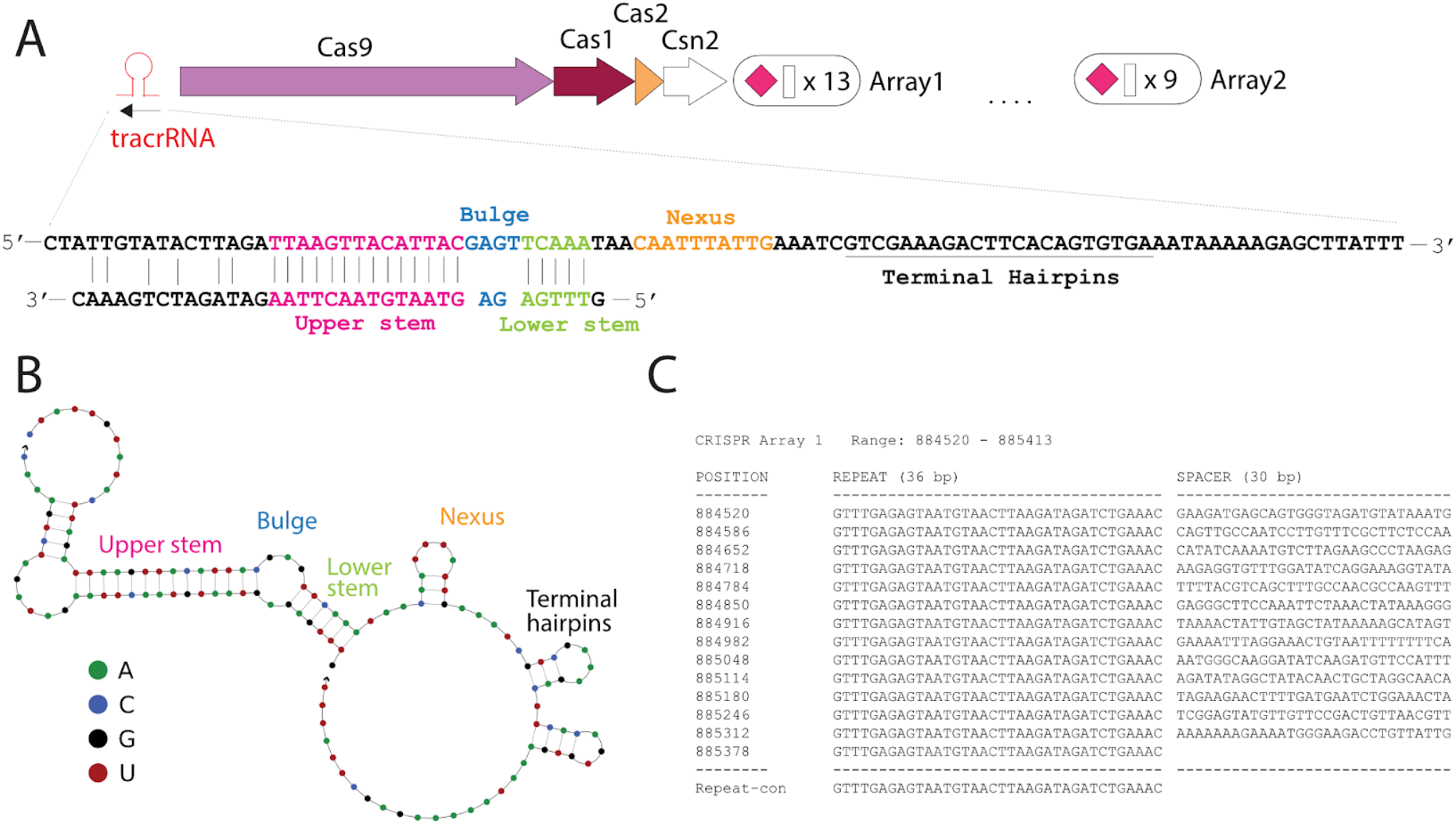
Characterization of CRISPR-Cas elements from *F. necrophorum* susbp. *funduliforme* 1_1_36S. **A**. Predicting the upstream *Trans-*activating CRISPR RNA (tracrRNA) of Cas9, coupled with the repeat region complementary to the tracrRNA. **B**. Prediction of the secondary structure of Cas9 tracrRNA. **C**. Overview of CRISPR repeats (36 bp) and spacers (30 bp) for the Cas9 locus of *F. necrophorum* subsp. *funduliforme* 1_1_36S.

The phylogenetic analysis of Cas9 from *F. necrophorum* subsp. *funduliforme* 1_1_36S revealed a conservation within the *Fusobacterium* genus **(Figure 2B)**. While the PAM sequence for *F. necrophorum* subsp. *funduliforme* 1_1_36S has not been experimentally characterized, the predicted PAM sequence for this Cas9 is NGG ^27^; the same as used for the commonly used *S. pyogenes* Cas9. By contrast, the predicted PAM sequence for the Type I-B CRISPR-Cas systems from *F. nucleatum* ATCC 23726 is YNW ^27^. Overall, a total of 3,427 spacers have been identified in Fusobacteria with only 629 matching viral, intergenic or ORFs ^28^. Utilizing viruSITE ^29^, the integrated database for viral genomics, most of the 13 spacers of the CRISPR array from *Fusobacterium* subsp. *funduliforme* 1_1_36S Cas9 were identified as a potentially part of *Synechococcus, Bacillus, Enterobacteria*, or *Aeromonas* phage genomes.

### Identification of the *F. necrophorum subsp. funduliforme* 1_1_36S and *F. ulcerans* 49185 Type IV CRISPR-Cas systems

Finally, Type VI CRISPR-Cas systems are divided into three types due to their low sequence similarity: VI-A, VI-B and VI-C. However, they all share the catalytic motif of the HEPN domain ^30^. The R-X4-H **(Figure S1)** motifs which are characteristic of HEPN-binding domains in both Cas13c that potentially mediate RNA cleavage is highlighted in **Figure 4** ^31^. *F. necrophorum* subsp. *funduliforme* 1_1_36S and *F. ulcerans* 49185 shared a 40.51% and 40.64% of identity when compared to the previously identified Cas13c from *F. perfoetens* ^*32*^. The Cas13c from *F. ulcerans* is composed of 1117 amino acids while the Cas13c from *F. necrophorum* is 1137 amino acids, which is consistent with the average size for Cas13c of approximately 1120 amino acids ^33^.

**Figure 4.**
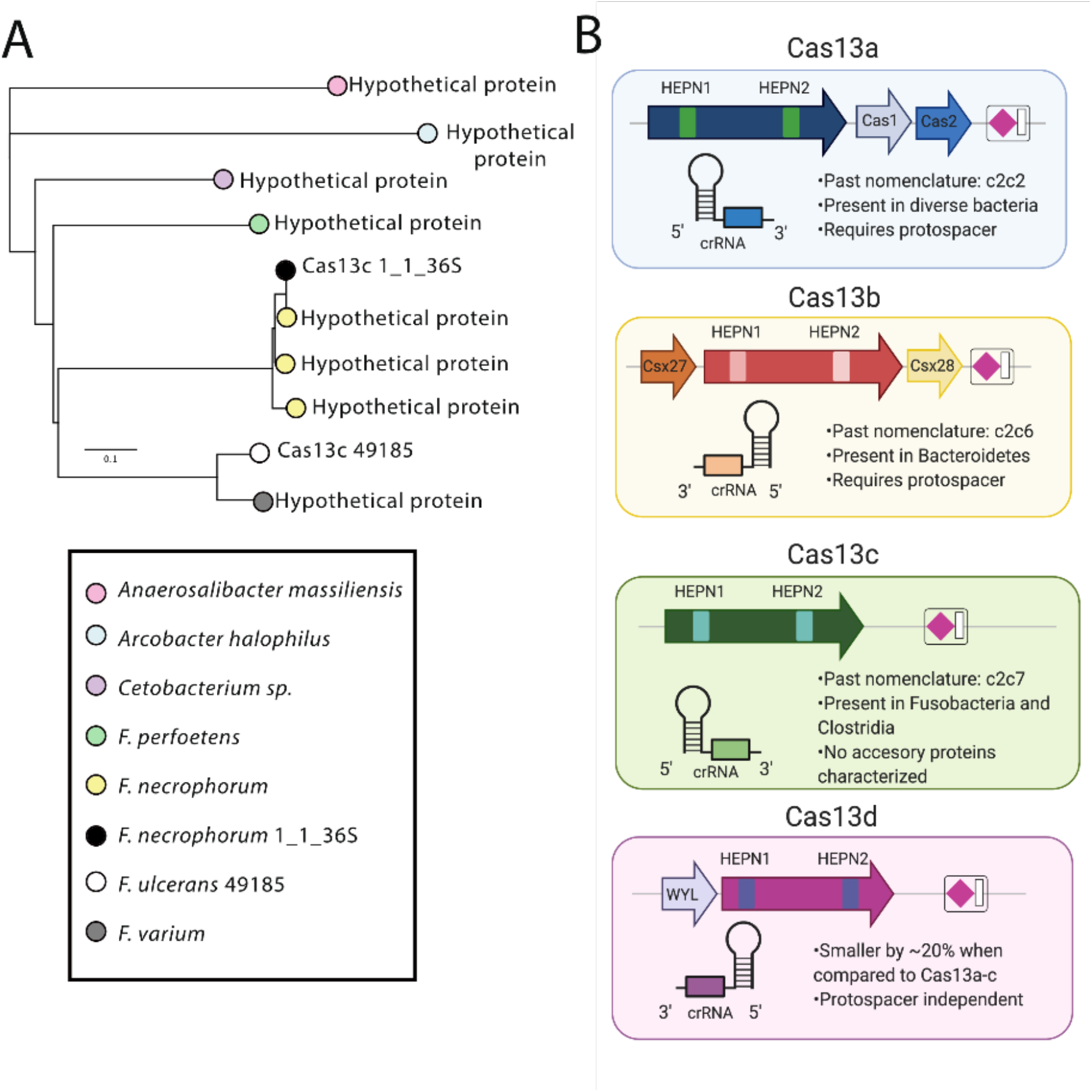
Analysis of Cas13c from a Type VI CRISPR-Cas system in *F. necrophorum* subsp. *funduliforme* 1_1_36S. **A**. SMART-BLAST phylogenetic analysis of Cas13c. **B**. domain organization of the subsets of Cas13 enzymes.

A phylogenetic analysis of bioinformatically predicted Cas13c uncovered the close phylogenetic relationship between Type VI effectors in *Fusobacterium* **(Figure 4A)**. Finally, when compared to other Type VI effectors Cas13c are the least studied and no accessory proteins have been identified like the other recently discovered Type VI systems.

## DISCUSSION

As CRISPR loci consist of host repeats and segments of foreign DNA extracted from invading organisms and mobile genetic elements, these repetitive stretches of DNA have proven difficult to resolve using short read DNA technologies that are commonly used to sequence bacterial genomes. The emergence of long-read DNA technologies from Oxford Nanopore and Pacific Biosciences has opened the door for the completion of fragmented genomes; many of which are likely incomplete due to the inability to resolve repeat regions. Previous attempts to determine the conformation, quantity, and diversity of CRISPR-Cas systems in *Fusobacterium* were exceptionally challenging because the complete genome sequencing information was incomplete, restricted, and incorrect. Complete and accurate genome sequences as the ones provided by FusoPortal are crucial for the analysis of these systems as well as the recombinant production of Cas proteins, or incorporation into a plasmid or chromosome; potentially leading to the expansion of *Fusobacterium* genetic toolkit ^24,34^.

A large percentage of bacteria harbor adaptive defense systems known as CRISPR-Cas systems, the analysis of CRISPR-Cas systems in *Fusobacterium* uncovered Class 1 and Class 2 systems. Class 2 systems are less common within prokaryotes when compared to Class 1 systems, at most 10% of CRISPR–Cas systems within sequenced genomes belong to Class 2 systems ^35^. For that reason, most of the identified CRISPR-Cas systems in *Fusobacterium* belong to Class 1 and interestingly, only *F. necrophorum* subsp. *funduliforme* 1_1_36S and *F. varium* 49185 encode for Class 2 systems which is similar to the CRISPR-Cas systems identified in *Leptotrichia*, which belongs to *Fusobacterium* genus, where they are unevenly distributed ^36^.

Type I CRISPR-Cas systems were found to be the most abundant in the eight *Fusobacterium* strains, which often coexist with Type III CRISPR-Cas systems as shown in *F. periodonticum* 2_1_31. It is suspected that Type I systems complement Type III systems functionally ^37^. Nonetheless, whereas Type I systems especially degrade dsDNA, Type III systems target ssDNA, dsDNA, and ssRNA ^37^. CRISPR mediates adaptive immunity by ensuing three steps. Particularly, most of *Fusobacterium* presented a complete operon of Type I-B CRISPR-Cas systems which is required to achieve adaptive immunity. First, the adaption step by Cas1, Cas2 and Cas4 requires the acquisition and inclusion of short segments of exogenous nucleotides into CRISPR loci; then crRNA-processing by Cas6 where the crRNA expression and maturation take place, and the last step is interference by Cas3, Cas5, Cas7 and Cas8 which recognize the crRNA and target nucleic acid ^38^. Generally, the Type I-B CRISPR-Cas system utilizes a cascade to target foreign DNA by base pairing CRISPR RNA (crRNA) to protospacers ^39^. In the case of *F. ulcerans* 49185 and *F. varium* 27725 which lack the signature *cas8* gene, the interference step could be defective as it has been demonstrated that Cas8 plays a key role in targeting cascade to invader DNA ^39^. Most Type I-B CRISPR-Cas systems in the analyzed eight strains were encoded downstream of the *cas2* gene and/or upstream of the *cas6* gene which is in accordance with a CRISPR-Cas system present in the anaerobic rod *Clostridium thermocellum* ^*40*^.

Type III-A CRISPR-Cas adaptive immune systems are composed of a protein-RNA complex 21. The complete loci in *F. periodonticum* 2_1_31 is constituted of five proteins and additional proteins that contain the Csm targeting complex Cas10/Csm1, Csm2–Csm5 and the Csm6 ribonuclease while *F. mortiferum* 9817 lacks Csm1 and Csm6 which could result in a non-functional CRISPR-Cas system. The role of these accessory genes may be related to modulation, complementation, or extension of these systems ^22,37,41^.

To determine how CRISPR-Cas systems will function the Type, Class and PAM recognition of the endonucleases needs to be characterized. For the Type II-A CRISPR-Cas system in *F. necrophorum* subsp. *necrophorum* 1_1_36S the PAM sequence was previously predicted as NGG and we further characterized its tracrRNA and spacer composition ^27^. This system could be utilized for genome modification in *Fusobacterium* strains **(Figure 5)**, however distinct strains of an organism encode as shown in this study, different elements to their CRISPR systems and consequently, may have unique requirements for genetic editing.

**Figure 5.**
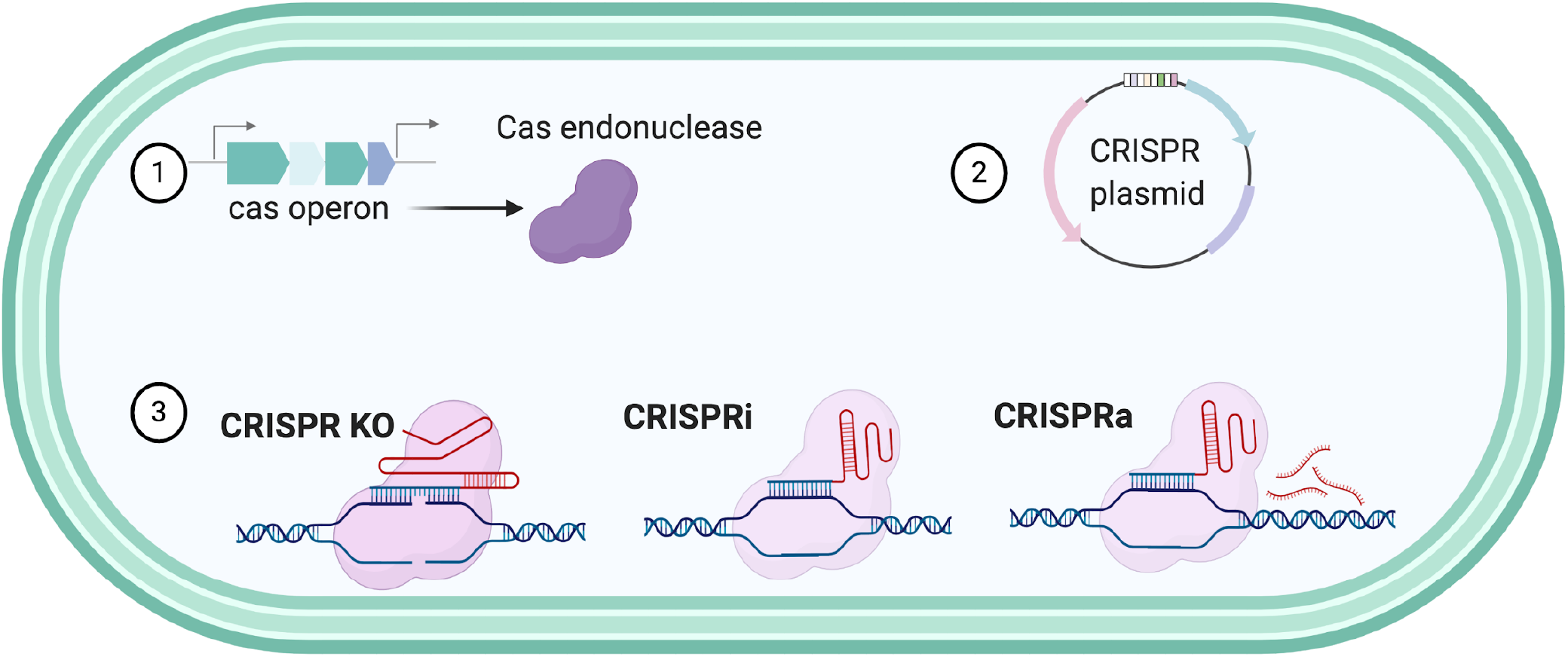
The use of native CRISPR-Cas systems as genetic tools could lead to the development of powerful tools. These tools could improve genome editing of *Fusobacterium* by creating deletion systems, characterization of essential genes through the CRISPRi, or the overexpression proteins using CRISPRa.

The most conserved trait in Type VI CRISPR-Cas is the HEPN domain ^30^. Interestingly enough, Type III systems possess RNase activity through the HEPN domain contained in the Csm6 and Csx1, although their role in this specific system remains to be studied ^42^. The first report of a Type VI system bioinformatically identified in *Fusobacterium* belonged to *F. perfoetens* ^*32*^, a strain that was found in children with diarrhea, nursing infants and in feces of a piglet. Additionally, CRISPR-Cas V1-C systems were only found in *Clostridium* and *Fusobacterium* ^*30*^.

The subtypes in the VI systems are based in the HEPN domains which are highly variable, the most widely studied being Type VI-A **(Figure 4)** ^11^. The HEPN domains are found frequently in various defense systems ^30^, in Cas13d proteins, this domain has been found to orchestrate CRISPR RNA maturation and target cleavage ^31^ which could suggest a potential role for the two HEPN domains identified in Cas13c from *F. necrophorum* subsp. *funduliforme* 1_1_36S and *F. ulcerans* 49185.

Additionally, CRISPR-Cas spacers represent a successful defense mechanism from past intruders, and the conserved spacers reflect the evolutionary phylogeny or conservation of species and strains ^36^. In the studied *Fusobacterium* strains, the spacers were not similar within them and the number variability suggests the biological heterogeneity of *Fusobacterium* species that encode a CRISPR-Cas system as a defense mechanism and consequently, for modulation of immunity ^36^. Higher number of spacers was observed in Type I-B which coincides with the fact that Type I-B, II-C and III-like CRISPR-Cas systems overall contain an increased number of spacers when compared to Type III-A, III-D, and VI-A.

In conclusion, the entirety of the *Fusobacterium* strains studied have endogenous systems that could be exploited as genetic tools for genomic engineering. The development of a CRISPR-Cas gene editing tool in *Fusobacterium* requires the description and detailed examination of the Type/Class of CRISPR array and identifying the PAM recognition sequence. CRISPR-Cas systems are a good candidate for genetic engineering since they depend on the endogenous systems that provide existing components and it is necessary to expand the genetic toolbox as *Fusobacterium* only has a few genetic systems described ^1,2^.

The utilization of bioinformatics has contributed to the continued discovery of novel CRISPR-Cas systems, such as Type V and Type VI CRISPR-Cas systems. As this bioinformatic study validates the presence of diverse CRISPR-Cas systems in *Fusobacterium*, we hypothesize that this technology could be used to more efficiently introduce gene interruptions, point mutations, and transcriptional control, through CRISPR interference (CRISPRi) and activation (CRISPRa) **(Figure 5)** ^43^. Furthermore, the CRISPR-Cas technologies and applications that have been developed are rapidly evolving and could enable countless novel applications ^32^, which is of substantial impact as most strains of *Fusobacterium* do not contain native plasmids and the need to create broad host plasmids that allow for pan-genetics through CRISPR technologies is a priority for the field.

## Acknowledgments

The authors would like to acknowledge Dr. Kira Marakova (NCBI) for the insightful information regarding the Cas13c and HEPN domain composition. This work was funded by the USDA National Institute of Food and Agriculture and the Virginia Tech College of Agriculture and Life Sciences. Select figures were created through Biorender.com.

## SUPPLEMENTAL FIGURES

**Figure S1.**
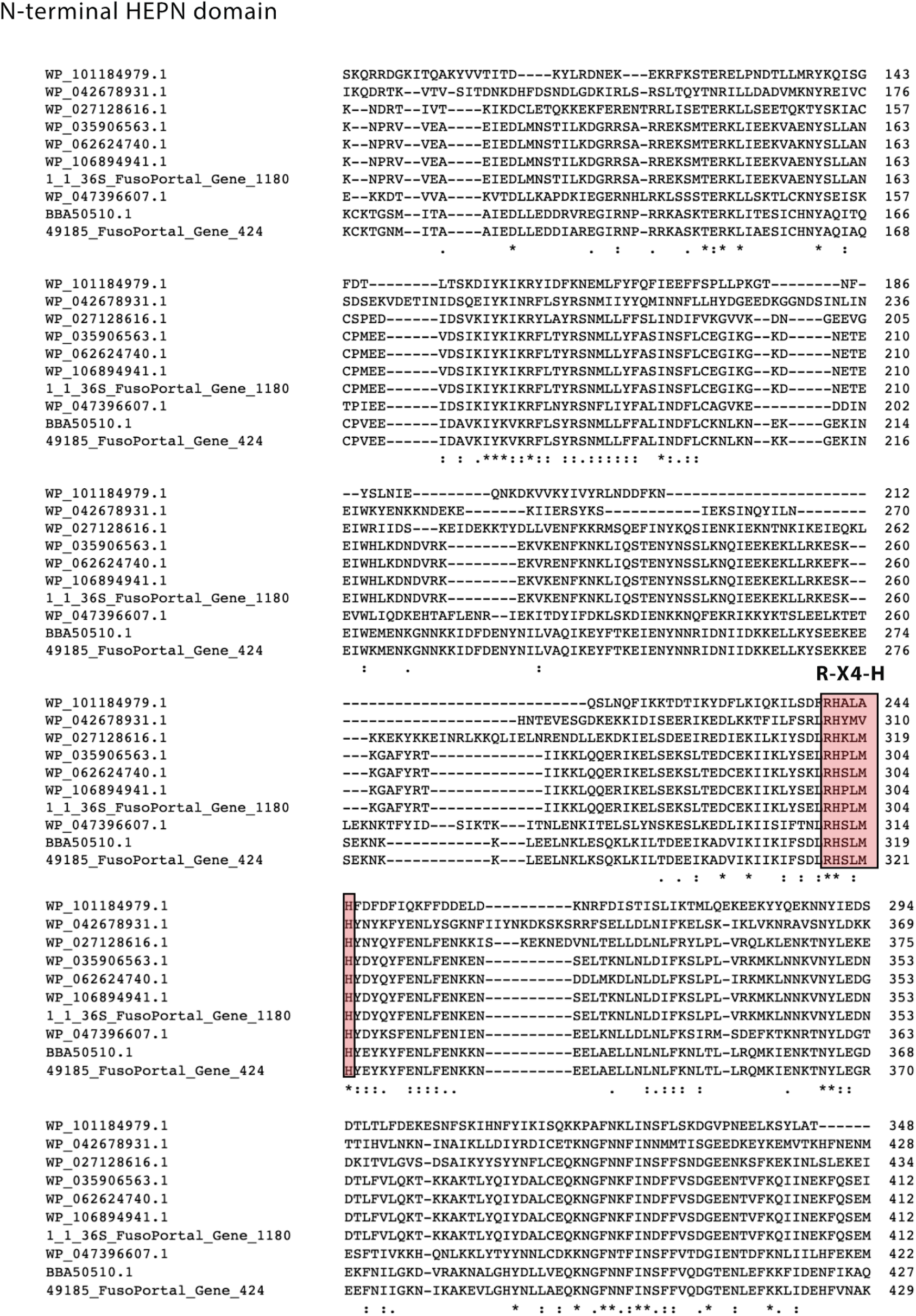
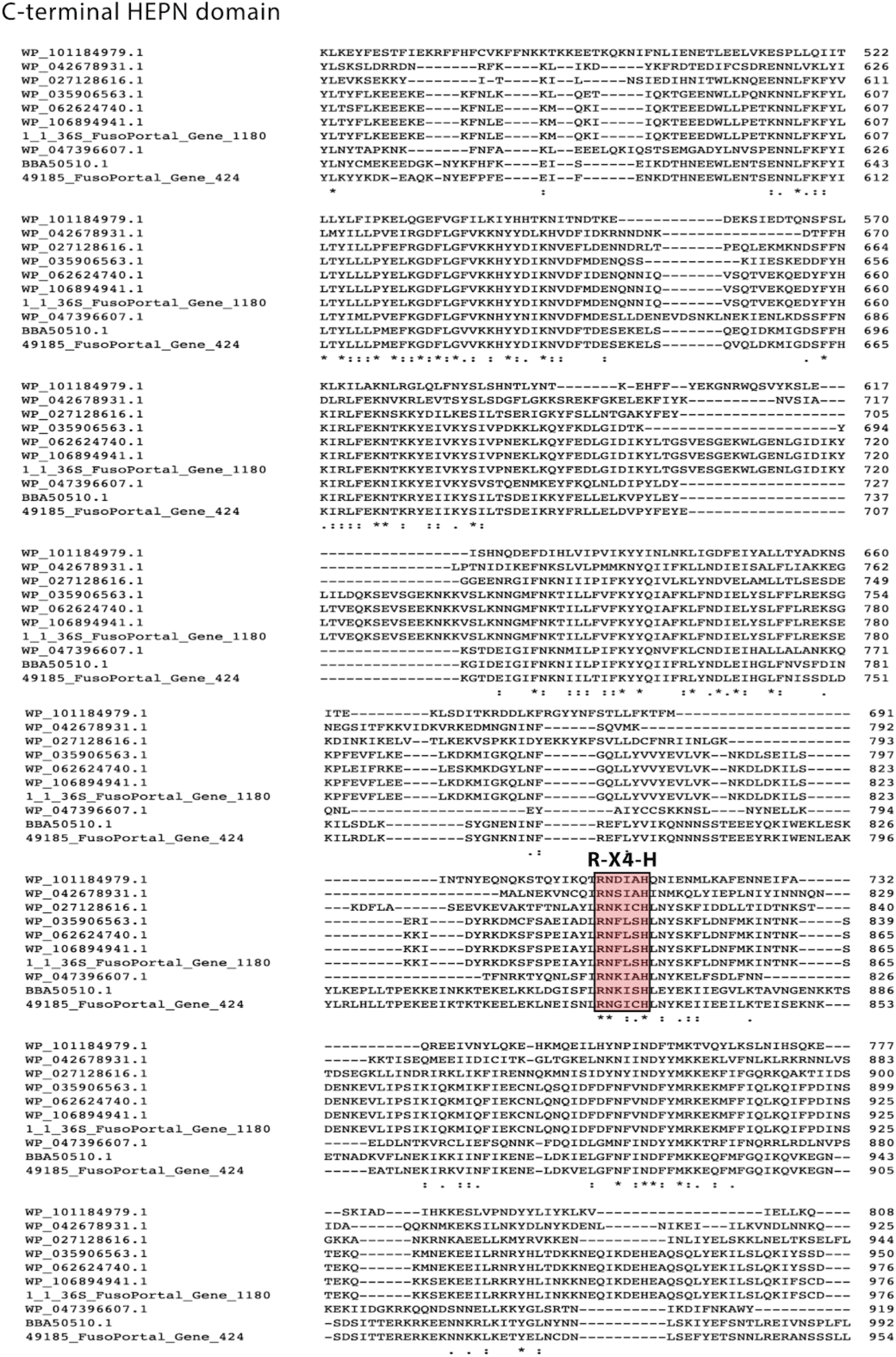
Multiple sequence alignment of Cas13c proteins. N-terminal and C-terminal HEPN domains are highlighted as well as the R-X4-H motif.

## SUPPLEMENTAL TABLES

**Table S1.** CRISPR-Cas system information of the eight analyzed *Fusobacterium* strains

| Species | Strain | CRISPR-Cas system Type | Subtype | Number of spacers | Conservation |
| --- | --- | --- | --- | --- | --- |
| *F. nucleatum* | 25586 | I | B1 | 15 | 0% |
| *F. nucleatum* | 23726 | I | B1 | 23 | 0% |
| *F. periodonticum* | 2_1_31 | I, III | B1, A2 | L1:42, L2:13, L3:13, L4:28 | L1:0%, L2:0%, L3:0%, L4:0% |
| *F. varium* | 27725 | I | B1 | L1: 13, L2:13, L3:24, L4:18 | L1: 9.75%, L2:0%, L3:0%, L4:2.04% |
| *F. ulcerans* | 49185 | I, VI | B1, C | L1:17, L2:23, L3:7 | L1:0%, L2:0%, L3:0% |
| *F. mortiferum* | 9817 | III | A1 | L1:4, L2:6 | L1:0%, L2:0% |
| *F. gonidiaformans* | 25563 | I | B1 | 83 | 0% |
| *F. necrophorum* | 1_1_36S | III, VI | A1, C | L1:13, L2:9 | L1:0%, L2:0% |

**Table S2.** CRISPR-Cas cluster elements in *F. nucleatum* ATCC 25586.

| Type | Cas proteins | Gene_FusoPortal_ID | GenBank_ID | Size (AA) |
| --- | --- | --- | --- | --- |
| I | Cas2 | 1711 | AVQ15518.1 | 92 |
| I | Cas1 | 1712 | AVQ15519.1 | 330 |
| I | Cas4 | 1713 | AVQ15520.1 | 164 |
| I | Cas3 | 1714 | AVQ15521.1 | 812 |
| I | Cas5 | 1715 | AVQ15522.1 | 366 |
| I | Cas7 | 1716 | AVQ15523.1 | 300 |
| I | Cas8a | 1717 | AVQ15524.1 | 517 |
| I | Cas6 | 1718 | AVQ15525.1 | 250 |

**Table S3.** CRISPR-Cas cluster elements in *F. nucleatum* ATCC 23726.

| Type | Cas proteins | Gene_FusoPortal_ID | GenBank_ID | Size (AA) |
| --- | --- | --- | --- | --- |
| I | Cas6 | 1183 | AVQ23192.1 | 250 |
| I | Cas8a | 1184 | AVQ23193.1 | 518 |
| I | Cas7 | 1185 | AVQ23194.1 | 300 |
| I | Cas5 | 1186 | AVQ23195.1 | 366 |
| I | Cas3 | 1187 | AVQ23196.1 | 812 |
| I | Cas4 | 1188 | AVQ23197.1 | 164 |
| I | Cas2 | 1190 | AVQ23199.1 | 92 |
| I | Cas1 | 1189 | AVQ23198.1 | 330 |

**Table S4.** CRISPR-Cas cluster elements in *F. ulcerans* 49185.

| Type | Cas proteins | Gene_FusoPortal_ID | GenBank_ID | Size (AA) |
| --- | --- | --- | --- | --- |
| I | Cas2 | 2775 | AVQ29172.1 | 94 |
| I | Cas1 | 2776 | AVQ29173.1 | 333 |
| I | Cas4 | 2777 | AVQ29174.1 | 160 |
| I | Cas3 | 2778 | AVQ29175.1 | 861 |
| I | Cas5 | 2779 | AVQ29176.1 | 251 |
| I | Cas7 | 2780 | AVQ29177.1 | 314 |
| I | Cas6 | 2782 | AVQ29179.1 | 213 |
| VI | Cas13c | 424 | AVQ26923.1 | 1117 |

**Table S5.** CRISPR-Cas cluster elements in *F. mortiferum* 9817.

| Type | Cas proteins | Gene_FusoPortal_ID | GenBank_ID | Size (AA) |
| --- | --- | --- | --- | --- |
| III | Cas6 | 2336 | AVQ19756.1 | 242 |
| III | Csm5 | 2337 | AVQ19757.1 | 383 |
| III | Csm4 | 2338 | AVQ19758.1 | 314 |
| III | Csm3 | 2339 | AVQ19759.1 | 231 |
| III | Csm2 | 2340 | AVQ19760.1 | 155 |
| III | Cas10 | 2341 | AVQ19761.1 | 797 |
| III | Cas2v7 | 2344 | AVQ19764.1 | 94 |
| III | Cas2III8 | 2345 | AVQ19765.1 | 96 |
| III | Cas1 | 2346 | AVQ19766.1 | 327 |

**Table S6.**
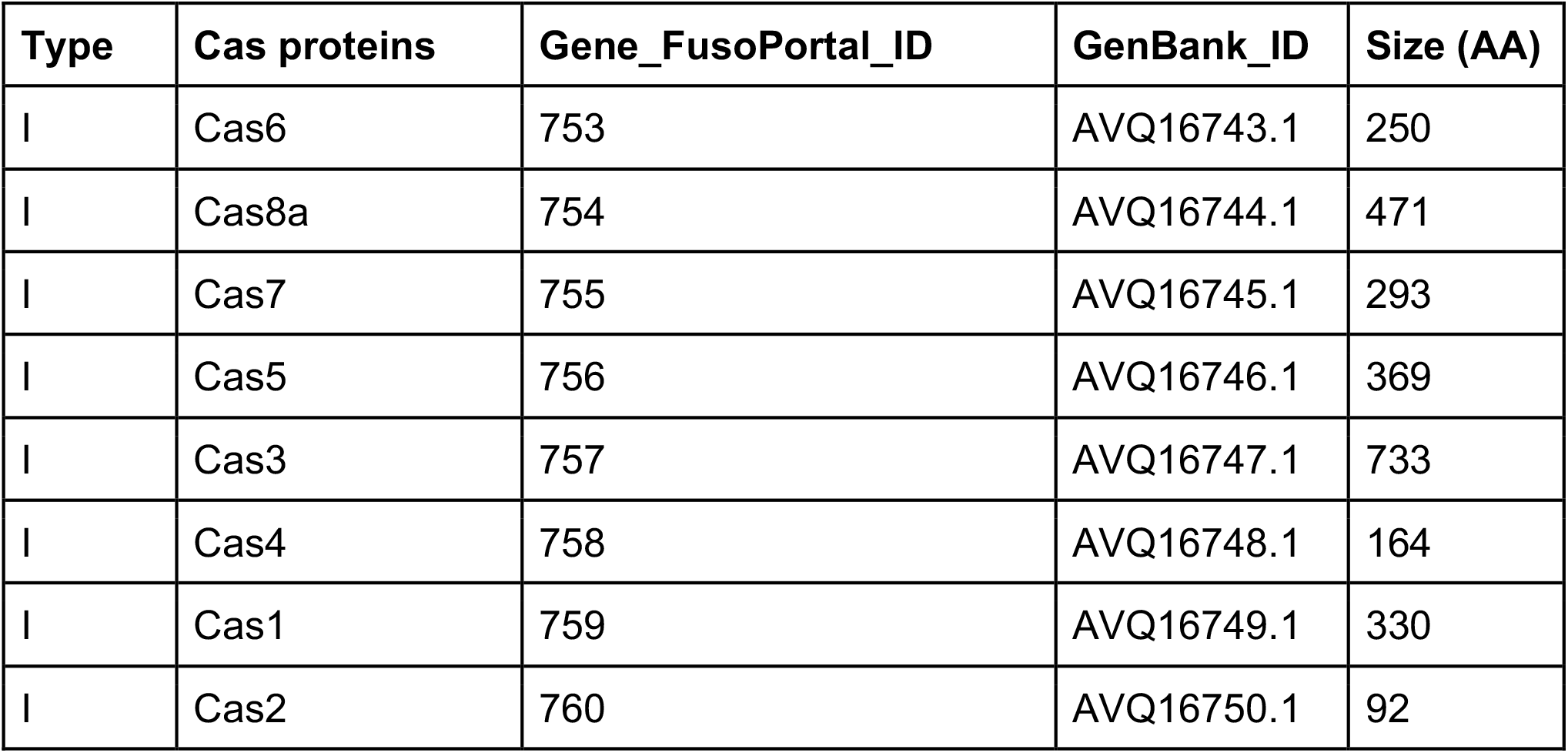
CRISPR-Cas cluster elements in *F. gonidiaformans 25563*.

| Type | Cas proteins | Gene_FusoPortal_ID | GenBank_ID | Size (AA) |
| --- | --- | --- | --- | --- |
| I | Cas6 | 753 | AVQ16743.1 | 250 |
| I | Cas8a | 754 | AVQ16744.1 | 471 |
| I | Cas7 | 755 | AVQ16745.1 | 293 |
| I | Cas5 | 756 | AVQ16746.1 | 369 |
| I | Cas3 | 757 | AVQ16747.1 | 733 |
| I | Cas4 | 758 | AVQ16748.1 | 164 |
| I | Cas1 | 759 | AVQ16749.1 | 330 |
| I | Cas2 | 760 | AVQ16750.1 | 92 |

**Table S7.**
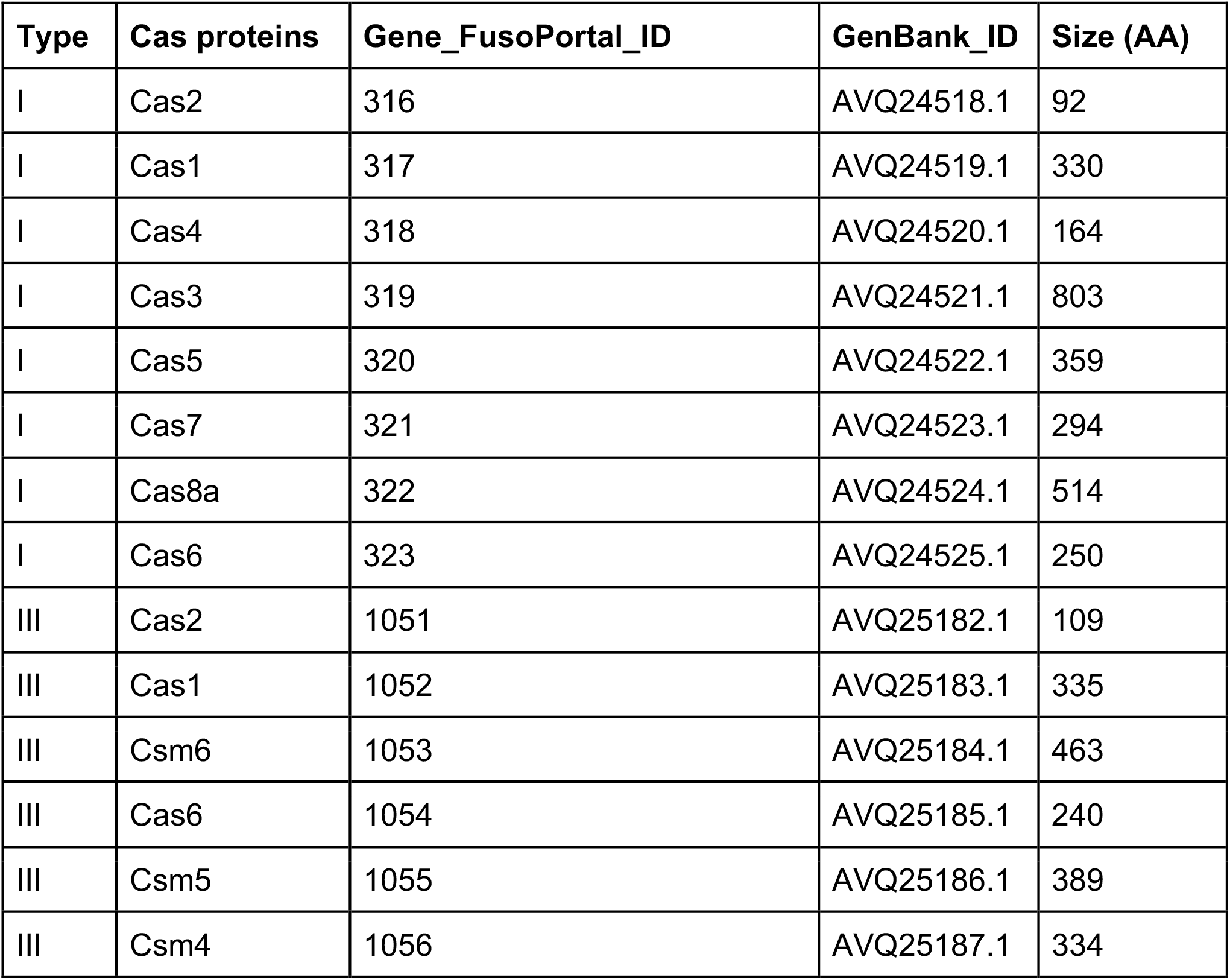
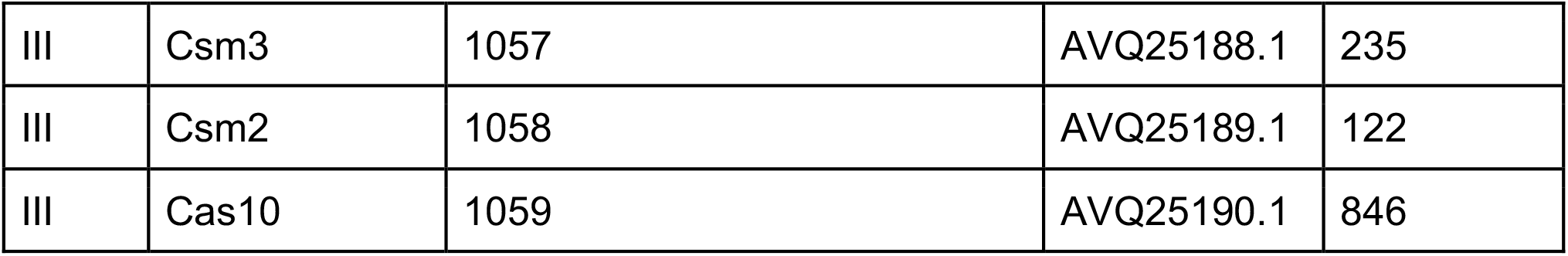
CRISPR-Cas cluster elements in *F. periodonticum* 2_1_31

| Type | Cas proteins | Gene_FusoPortal_ID | GenBank_ID | Size (AA) |
| --- | --- | --- | --- | --- |
| I | Cas2 | 316 | AVQ24518.1 | 92 |
| I | Cas1 | 317 | AVQ24519.1 | 330 |
| I | Cas4 | 318 | AVQ24520.1 | 164 |
| I | Cas3 | 319 | AVQ24521.1 | 803 |
| I | Cas5 | 320 | AVQ24522.1 | 359 |
| I | Cas7 | 321 | AVQ24523.1 | 294 |
| I | Cas8a | 322 | AVQ24524.1 | 514 |
| I | Cas6 | 323 | AVQ24525.1 | 250 |
| III | Cas2 | 1051 | AVQ25182.1 | 109 |
| III | Cas1 | 1052 | AVQ25183.1 | 335 |
| III | Csm6 | 1053 | AVQ25184.1 | 463 |
| III | Cas6 | 1054 | AVQ25185.1 | 240 |
| III | Csm5 | 1055 | AVQ25186.1 | 389 |
| III | Csm4 | 1056 | AVQ25187.1 | 334 |
| III | Csm3 | 1057 | AVQ25188.1 | 235 |
| III | Csm2 | 1058 | AVQ25189.1 | 122 |
| III | Cas10 | 1059 | AVQ25190.1 | 846 |

**Table S8.** CRISPR-Cas cluster elements in *F. necrophorum* subsp. *funduliforme* 1_1_36S.

| Type | Cas proteins | Gene_FusoPortal_ID | GenBank_ID | Size (AA) |
| --- | --- | --- | --- | --- |
| II | Cas9 | 859 | AVQ20950.1 | 1337 |
| II | Cas1 | 860 | AVQ20951.1 | 290 |
| II | Cas2 | 861 | AVQ20952.1 | 106 |
| II | Csn2 | 862 | AVQ20953.1 | 226 |
| VI | Cas13c | 1180 | AVQ21256.1 | 1137 |

**Table S9.** CRISPR-Cas cluster elements in *F. varium* 27725.

| Type | Cas proteins | Gene_FusoPortal_ID | GenBank_ID | Size (AA) |
| --- | --- | --- | --- | --- |
| I | Cas1 | 2395 | AVQ31955.1 | 333 |
| I | Cas2 | 2396 | AVQ31956.1 | 94 |
| I | Cas3 | 2393 | AVQ31953.1 | 858 |
| I | Cas4 | 2394 | AVQ31954.1 | 160 |
| I | Cas5 | 2392 | AVQ31952.1 | 250 |
| I | Cas6 | 2389 | AVQ31949.1 | 231 |
| I | Cas7 | 2391 | AVQ31951.1 | 314 |

